# A Mechanistic Approach to Optimize Combination Antibiotic Therapy

**DOI:** 10.1101/2024.06.10.598196

**Authors:** F. Clarelli, P. O. Ankomah, H. Weiss, J. M. Conway, G. Forsdahl, P. Abel zur Wiesch

## Abstract

Antimicrobial resistance is one of the most significant healthcare challenges of our times. Multidrug or combination therapies are sometimes required to treat severe infections; for example, the current protocols to treat pulmonary tuberculosis combine four antibiotics. However, combination therapy is usually based on lengthy empirical trials and it is difficult to predict its efficacy. We propose a new tool to identify antibiotic synergy or antagonism and optimize combination therapies. Our model explicitly incorporates the mechanisms of individual drug action and estimates their combined effect using a mechanistic approach. By quantifying the impact on growth and death of a bacterial population, we can identify optimal combinations of multiple drugs. Our approach also allows for the investigation of the drugs’ actions and the testing of theoretical hypotheses.

We demonstrate the utility of this tool with in vitro *Escherichia coli* data using a combination of ampicillin and ciprofloxacin. In contrast to previous interpretations, our model finds a slight synergy between the antibiotics. Our mechanistic model allows investigating possible causes of the synergy.

## 1. Introduction

Antimicrobial resistance (AMR) is a severe global health challenge [1]. A 2019 World Health Organization (WHO) report estimates that AMR causes 700,000 deaths per year, including 230,000 deaths from multidrug-resistant tuberculosis (MDR-TB) [2]. The report claims that, if we do not take action soon [3, 4], the annual death toll could reach ten million by 2050.

New antibiotics require years of expensive development and testing [5]. Researchers are also exploring new strategies that optimize the action of the antibiotic on bacterial population growth, while also minimizing the risk of selecting resistant strains [6, 7].

Physicians use multidrug or combination therapies to treat some bacterial infections (e.g. pulmonary tuberculosis), and provide empirical treatment for some critically ill (e.g. septic) patients [8]. Multidrug therapy is sometimes considerably more effective than monotherapy at clearing an infection, but the risk of selecting resistant strains may be higher if not carefully applied [9]. Models, especially mechanistic models [10], may provide a helpful tool to balance these considerations, and to suggest optimal treatments to improve the efficacy of the treatments and reduce the spread of AMR.

Several approaches are used to characterize the effects of a drug combination on bacteria. A detailed review is in [11], and additional works describing the drug combination and how to combine the two drugs (e.g. Bliss independence and Loewe additivity) are in [12, 13]. The principal methods used to describe the drug effect rely on the Emax or Hill curves and are sometimes based on a multi-hit model [13].

Drug-drug interaction is a critical problem in pharmacology (e.g. see EMA guidelines [14]); although pharmacokinetic (PK) interactions are well studied, the study of pharmacodynamic (PD) interactions often focuses on descriptive methods.

In this work, we propose a new mechanistic (bottom-up) approach to describe the dose-response effects of a drug combination. We consider the action of antibiotics via target binding: the more drug molecules that bind to their targets, the greater the damage to the bacteria. This bridge between the intracellular action of antibiotics and the bacterial population’s growth (and death) differs from other pharmacodynamic approaches [15-18]. In other words, we introduce a model that describes the action of drugs in killing bacteria by connecting the number of bound targets with the killing or inhibition of replication of bacteria. The number of bound targets is a function of the drug concentration; in this way, we obtain a mechanistic exposure-response relationship. Previous models have only connected drug-target binding to bacterial death, but not bacterial replication. Such an approach is appropriate for antimicrobial peptides [13] and purely bactericidal antibiotics, but not for antibiotics that inhibit replication, i.e. those with at least partial bacteriostatic action.

We postulate a different form of characterizing the effects of independent drug combinations by combining drug effects linearly on a mechanistic basis with no interaction. We use time-kill curve experiments to calibrate growth and death rates of bacteria under different concentrations of drugs (in monotherapy). After the calibration, we can simulate the growth of bacteria populations under any constant or time-dependent drug combination, and estimate the most efficient combination. This new approach allows us to detect and quantify drug synergies.

In the following sections, we introduce the model and its effects for a two-drug combination. We subsequently show how to extend it to four drugs. We test our model with a combination of ciprofloxacin and ampicillin to treat *E. coli* bacteria (data published in [19]). We observe a synergistic effect of the two drugs, especially at high concentrations, and then investigate the possible causes of the synergy. Finally, we introduce a theoretical resistant strain example to show how our model can predict the selection pressure for the resistant strain under different drug combinations, and how a mutated strain can change the efficacy of the treatment.

## 2. Methods

We design a mathematical model to describe the action of drugs on bacteria. Our model describes the dynamics of bound targets inside bacteria using the association and dissociation rates of antibiotics with their targets. There are two overall effects of the drugs on the cell population: reduction of growth rate (bacteriostatic action) and increase in death rate (bactericidal action). These two effects are functions of the number of bound targets caused by the drugs administered. When no target is bound by drug molecules, bacteria have maximum growth and natural death rates.

The calibration of the model establishes the relationship between the number of bound targets and the growth and death rate of the bacterial population. After model calibration, we can simulate the bacterial population’s response to any amount of drug (exposure-response).

Next, we extend this concept to multiple drugs, adding a new target for each new drug (if it binds with a different target). In this way, we can evaluate the effect of multiple target binding, and ascertain which combinations of antibiotics can be more effective.

### 2.1 Model description

We consider a bacterial population, B(t), where each bacterium has two different targets. For the sake of simplicity, we assume that each bacterium has the same mean number of free and bound targets. A list of the variables used in the equations is presented in Table 1.

**Table 1.** In the table, we introduce all the variables used in the equations. The units are numbers of elements.

| Description | Drug 1 | Drug 2 |
| --- | --- | --- |
| Number of drug molecules | $A_1$ | $A_2$ |
| Free targets per bacterium | $\tau_1(t)$ | $\tau_2(t)$ |
| Bound targets per bacterium | $\alpha\tau_1(t)$ | $\alpha\tau_2(t)$ |
| Number of targets per bacterium (constant) | $\theta_1 = \tau_1(t) + \alpha\tau_1(t)$ | $\theta_2 = \tau_2(t) + \alpha\tau_2(t)$ |
| Total number of free targets (in population) | $T_1(t) = B(t) \cdot \tau_1(t)$ | $T_2(t) = B(t) \cdot \tau_2(t)$ |
| Total number of bound targets (in population) | $AT_1(t) = B(t) \cdot \alpha\tau_1(t)$ | $AT_2(t) = B(t) \cdot \alpha\tau_2(t)$ |
| Total number of targets (in population) | $\gamma_1(t) = B(t) \cdot \theta_1$ | $\gamma_2(t) = B(t) \cdot \theta_2$ |
| Number of bacteria | $B(t)$ | |

#### 2.1.1 Association and dissociation rate

We now introduce how the association and dissociation rates affect the dynamics of bound targets. We indicate with *A*_*i*_ the number of molecules of the two drugs (i=1,2). They bind to their targets with a rate 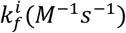 and unbind with a rate 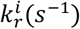. As we use different units in our equations, we prefer to rescale the rate 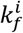 to 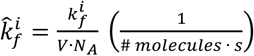, where *N*_4_ is the Avogadro number, M is the molarity (mol/L), and *V* is the volume where we have a homogeneous distribution of the drug molecules and targets. Considering that i indicates the drug, the contribution of the association and dissociation terms is:

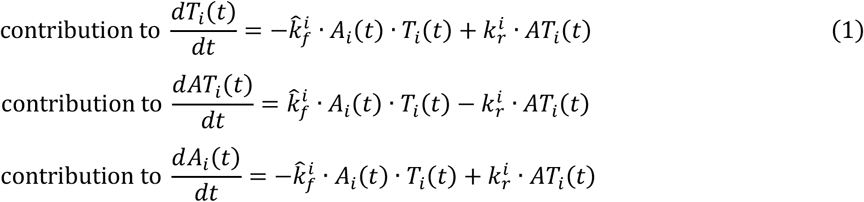

#### 2.1.2 Replication rate

We can assume that the cell population grows exponentially for some time, with a rate *r* when no drugs are applied [20], thus

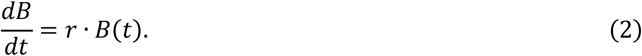

The drug molecules affect the growth rate of bacteria. We present an example to illustrate what happens to the bound targets during bacteria replication. If we consider only the duplication of a bacterium with four bound targets and four free targets (Figure 1), we still have four bound targets in total after the duplication – two per bacterium. The total number of bound targets does not change in the duplication of bacteria. We can therefore assume that the total number of bound targets (in the population) remains constant: i.e. *AT*_*i*_ = *constant*, which implies that the new targets arising during the replication are all free targets (see the number of red arches in Figure 1). The bound targets vary when we add the association term, the dissociation term and the death term to the equations.

**Figure 1.**
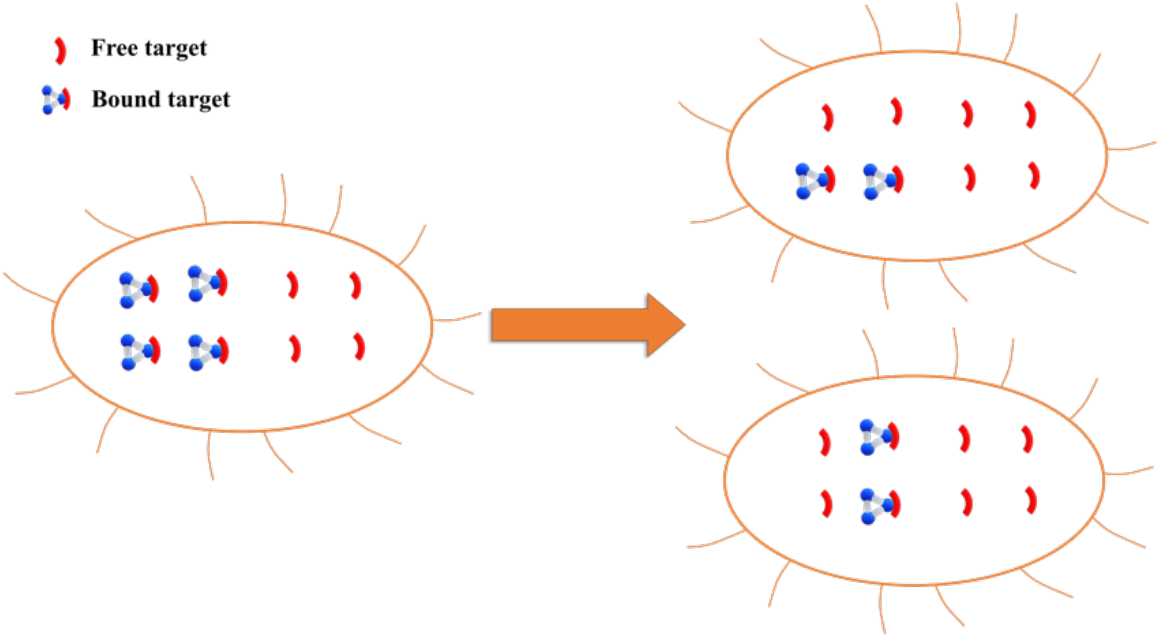
Duplication of bacteria. The number of bound targets remains constant during the duplication of a bacterium with four bound targets. For this reason, we have two daughter cells with two bound targets per bacterium. increases, the replication rate decreases until it can become zero (complete growth inhibition). We can estimate the growth rate r = r(AT_1_, AT_2_) by calibrating the model with experimental time-kill curves that describe the bacterial growth under different drug concentrations.

We can write the cell population growth rate as a function of bound targets (bacteriostatic action) *r* = *r*(*AT*_1_, *AT*_2_). With no drug, the replication rate has its maximum value *r*(0) = *r*_*max*_. When the number of bound targets

Under these assumptions, we can rewrite the growth of bacteria *B(t)* as the growth of free and bound targets *T*(*t*) and *AT*(*t*). We can rewrite equation (2) yielding contribution of bacterial replication to *T*_*i*_ and *AT*,

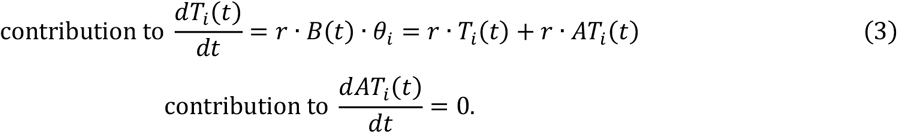

#### 2.1.3 Death rate

The number of bound targets also affects the death rate of bacteria δ. With no drugs, the death rate is zero or equal to the natural death rate of the bacteria. When the number of bound targets increases with higher drug concentration, the death rate increases until reaching its maximum value δ_*max*_. We can estimate the death rate function by calibrating the model with experimental time-kill curves. Representing the death rate as δ(*AT*_*i*_), we have

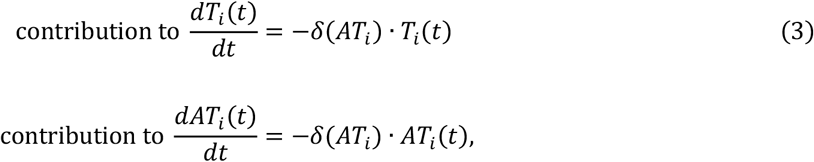

where i=1,2.

We now combine all the four terms: association rate, dissociation rate, growth rate and death rate.

#### 2.1.4 Full system

The model for two drugs is

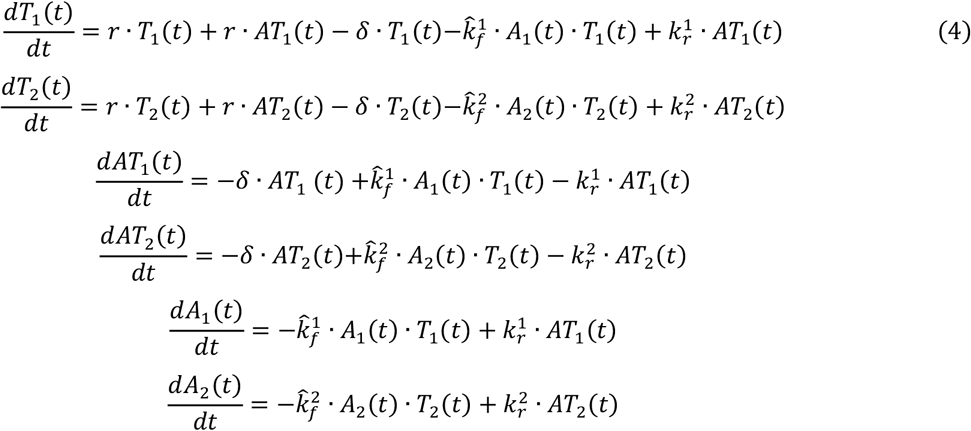

Based on the assumption that all the targets are initially free and the number of bound targets is zero, we use the following initial conditions: *T*_*i*_(0) = *B*(0) ⋅θ_*i*_, *AT*_*i*_ (0) = 0, and *A*_*i*_ (0) = initial # of drug molecules, for i=1,2.

We solve the ordinary differential equation system numerically. In the following sections, we use the minimum squared error to obtain the best fit of this model with the experimental data. To calibrate the model, we use particle swarm optimization (PSO), which is a global optimization method. We use Matlab R2021 (MathWorks) to perform the numerical simulations.

### 2.2 Drug combination

We introduce a mechanistic model where the number of drug molecules affects the bacteria’s growth and death rates. In the case of two drugs, we have *r*_1_(*AT*_1_), *r*_2_(*AT*_2_), δ_1_(*AT*_1_) and δ_2_(*AT*_2_), where *AT*_1_ and *AT*_2_ are the numbers of bound targets for the first and the second drug, respectively, as introduced in Table 1.

#### 2.2.1 Effect on the growth rate

We define the effect of drug molecules on the growth rate as the difference between the optimal growth rate and the actual growth rate in the presence of drugs: 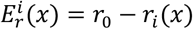. A visual example is presented in Figure 2A, where *r*_0_ is the growth rate with no drugs.

**Figure 2.**
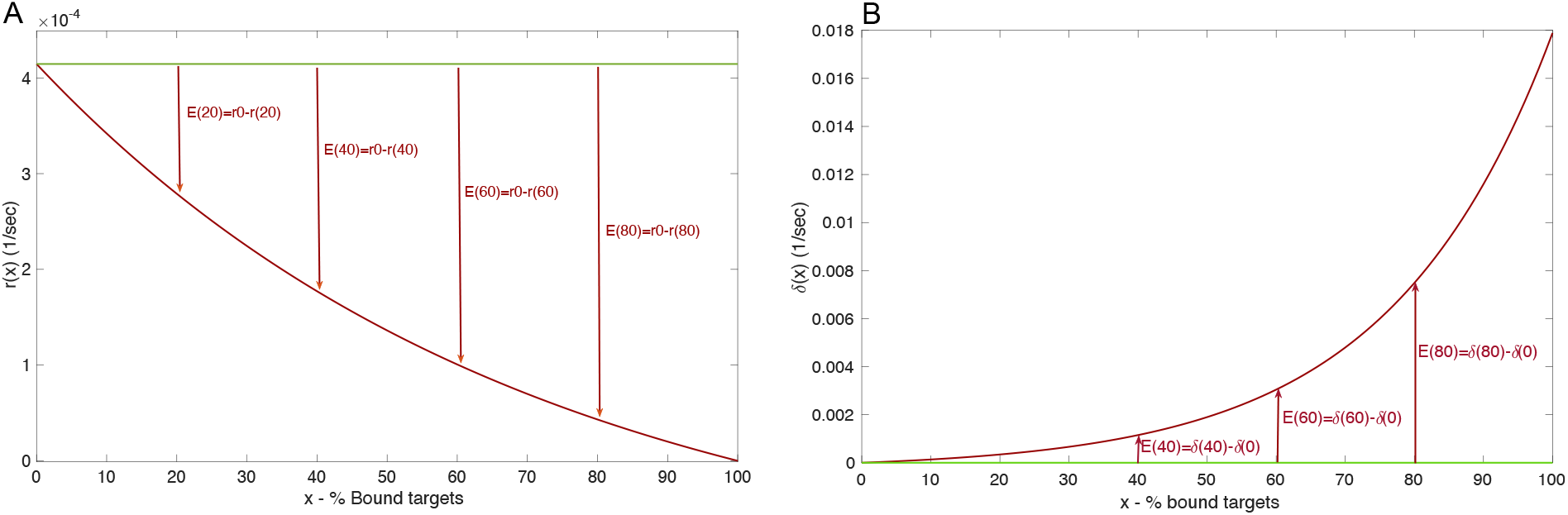
**A**: Vertical red arrows represent the amplitude of the drug effect on the growth rate (bacteriostatic effect). The green line represents the optimal growth rate, and the red line is the growth rate reduced by antibiotics. **B**: Vertical red arrows describe the effect of a drug on the death rate (bactericidal). The green line represents the natural death rate, and the red line the increased death rate.

#### 2.2.2 Effect on the death rate

The difference between the actual death rate δ_*i*_(*x*), in the presence of drug molecules, and the natural death rate δ(0) gives us the effect 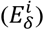 of the drug molecules i on the death of bacteria: 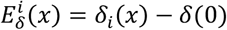. If we neglect the natural death rate of bacteria, we have 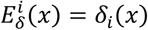. A visual example is presented in Figure 2B, where δ(0) is the natural death rate.

#### 2.2.3 Independent combination

Several articles describe the effect of drug combinations; see, for example, [13, 21]. These are often based on a more phenomenological approach. The preliminary step in our analysis is to define a ‘frame of reference’ – e.g. the independent combination of the growth rate effects. This is the linear combination of the two single effects: 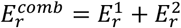. Thus, the growth rate corresponding to the combination of two drugs is

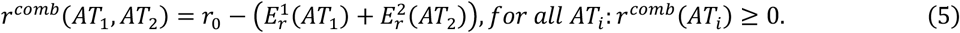

Similarly, an independent combination of the effects on the death rate is 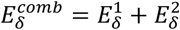, and the death rate corresponding to the combination of two drugs is

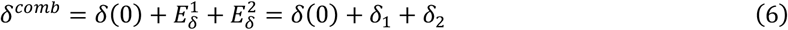

Here we define a linear independent combination of effects.

#### 2.2.4 Synergistic and antagonistic effects

We define a local (i.e. dependent on the drug concentration) synergy for the growth rate when *r* < *r*^*comb*^, and a local antagonism if *r* > *r*^*comb*^. Similarly, we define a local synergy for the death rate when δ > δ^*comb*^, and a local antagonism when δ < δ^*comb*^. In our model, drug-drug interactions are described using the parameter α. In our example, we focus on the interaction of ampicillin and ciprofloxacin. Ampicillin is.commonly thought to be a purely cidal antibiotic, i.e., has no effect on bacterial growth. Therefore, we here only focus on the interaction of antibiotic induced death rates. The expected death rate δ^*comb*^_*e*_is calculated as described above, and compared to the fit, resulting in a new death rate δ_*comb*_ = α · (δ_*cipro*_ + δ_*amp*_). Values of α > 1 indicate synergy, values of α < 1 antagonism. While our hypothesized formulation to model synergy/antagonism is simple, we lack the data to motivate more complex functional forms, and this formulation explains the data and provides biological insights (a mechanistic discussion will follow further down). As denoted in table 1 and below in the results section, α can also be used to modify the number of bound target molecules or the effective drug concentration inside the bacteria. While our hypothesized formulation to model synergy/antagonism is simple, we lack the data to motivate more complex functional forms, and this formulation explains the data and provides biological insights.

### 2.3 Four drug extension

Motivated by the current treatment of pulmonary tuberculosis, we now extend our previously discussed model to four antibiotics (with four different targets). Several new drugs (e.g. linezolid, moxifloxacin, pretomanid, bedaquiline) are entering the TB treatment space, particularly for use in treating MDR-TB [22, 23], and a mechanistic framework to assess their potential effect in combination has not been forthcoming. We use the same structure introduced above and the variables summarised in Table 1, where the index i now ranges from 1 to 4. We have a population of bacteria B with four different targets. As previously assumed, θ_*i*_ represents the total number of targets per bacterium, 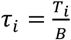 is the average number of free targets per bacterium, and 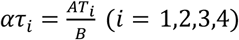 1,2,3,4) is the average number of bound targets per bacterium. The whole system with four drugs is

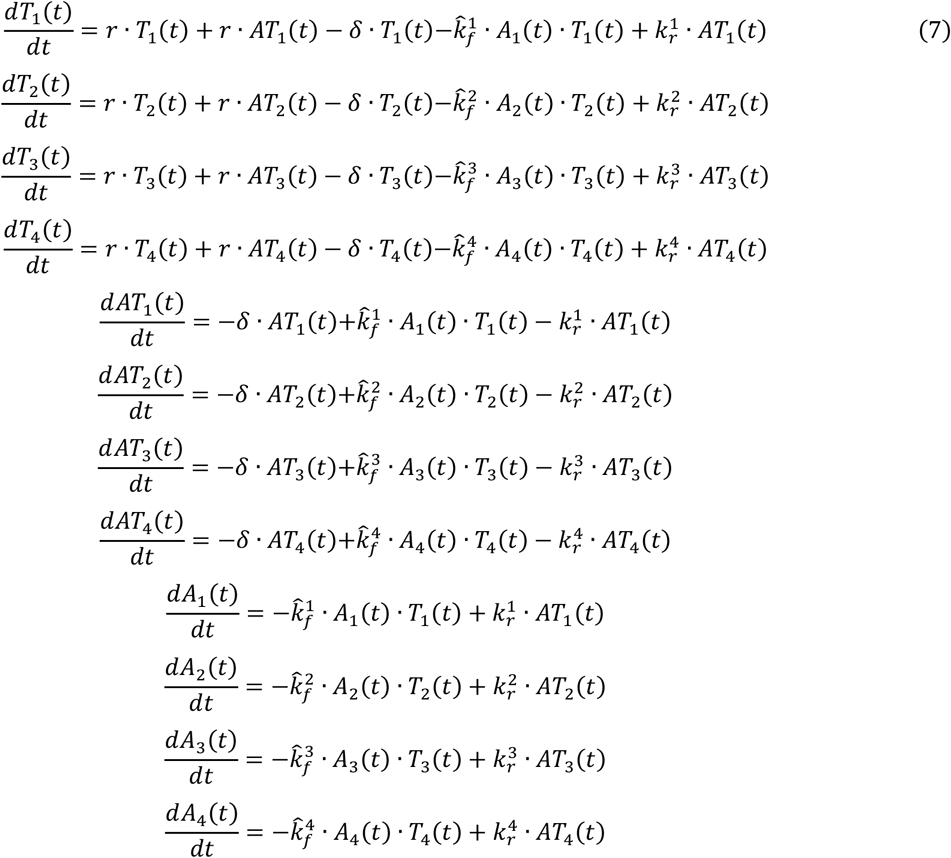

## 3. Results

We test our two drug model with experiments where *E. coli* was exposed to combinations of ampicillin and ciprofloxacin (so called time-kill curves, published in [19]). These experiments are clinically relevant because clinicians sometimes treat gram-negative infections using a combination of beta-lactams and fluoroquinolones, with some studies demonstrating a clinical advantage to the combination [24]. In these experiments, *E. coli* were exposed to rising concentrations of the two antibiotics and the size of the bacterial population was followed over time. This is a measure of drug efficacy, and generally speaking, the rate with which the bacterial population declines, increases with the antibiotic concentration.

We have previously used a similar model for a single drug for describing the effects of ampicillin and ciprofloxacin on *E. coli* individually [17]. However, these results were obtained with *E. coli* strain BW25113, while the results with both drugs combined were obtained with *E. coli* strain CAB1. While we assume that drug-target affinity is comparable by taking these values from the same literature, we assume that the effects of successive target binding on bacterial growth and death are different. We therefore first need to fit our model to *E. coli* CAB1 exposed to ampicillin and ciprofloxacin individually (see supplementary material). We then simulate the effects of an independent drug combination (as introduced above) on the bacterial population and compare the simulations with the experimental data of the corresponding drug combinations.

### 3.1 Combination ciprofloxacin-ampicillin

We assume the same association and dissociation rates used to model bacterial populations exposed to single drugs to model the drug combination. In addition, ampicillin and ciprofloxacin concentrations in the experimental work were selected as corresponding multiples of the in vitro MIC for each individual antibiotic. For this reason, the ratio between the concentrations of the two drugs is constant (115.66) [19]. The underlying experimental data are shown in Figure 3.

**Figure 3.**
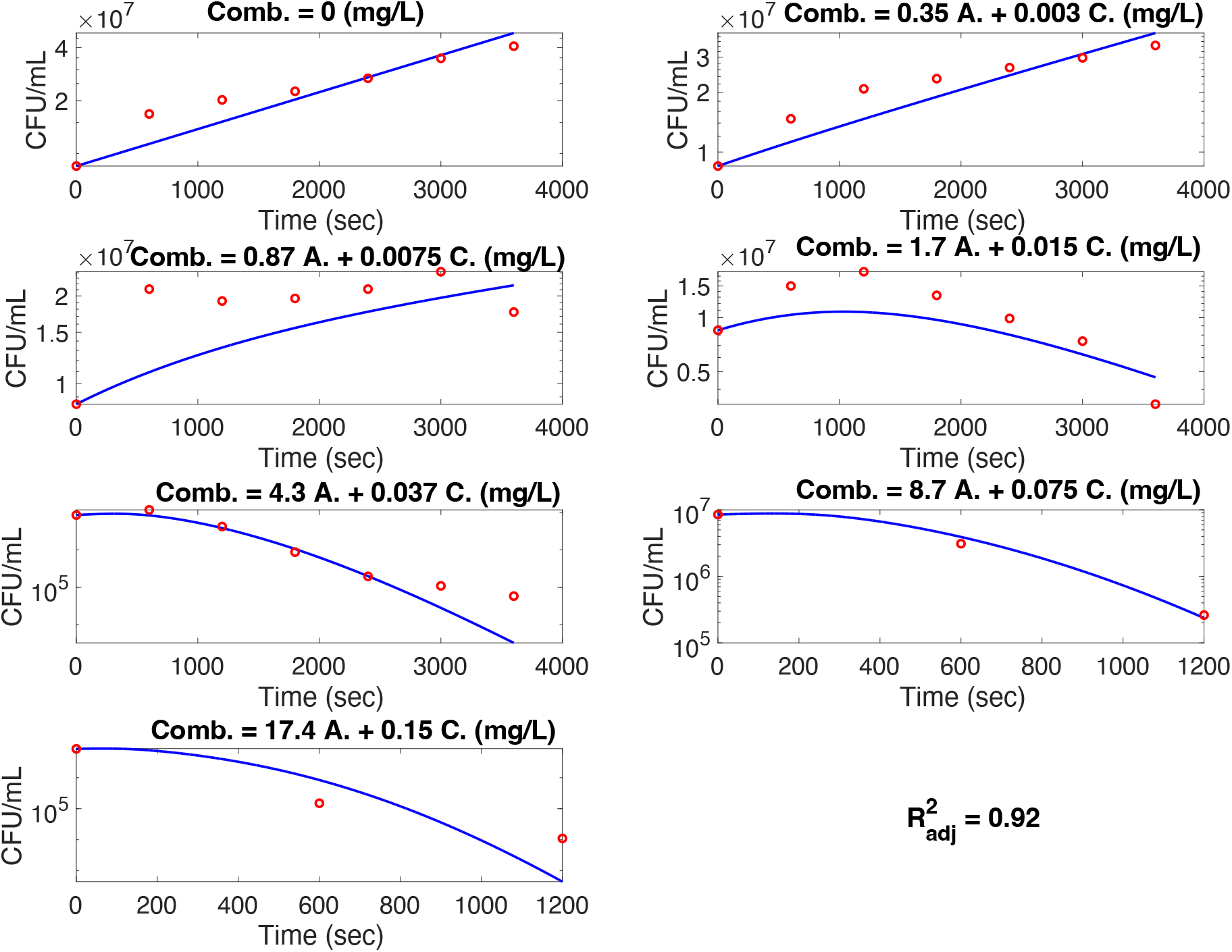
The best fit of the ampicillin-ciprofloxacin combination. In the last two concentrations, the third figure of the right column and the fourth figure of the left column, we neglect the experimental points where persistence appeared. Therefore, we only include the first three experimental points (red stars).

#### 3.1.1 Growth rate

The growth rate of bacteria *r* with two drugs is equal to the growth rate of ciprofloxacin alone because we neglect the bacteriostatic action of ampicillin.

#### 3.1.2 Death rate

As introduced in the methods section, the resultant death rate is the linear combination of the ciprofloxacin and ampicillin death rate functions: δ_*comb*_ = α · (δ_*cipro*._ + δ_*amp*_), where the coefficient α determines whether the action of the two drugs is synergistic (α > 1), or antagonistic (α < 1). If α = 1, we define δ_*indep*_ = (δ_*cipro*_ + δ_*amp*_) as a linear independent combination of the two effects. We can empirically calculate the factor α by fitting our model with seven experimental time-kill curves obtained with the drug combination published in [19]. These combinations have the same ampicillin and ciprofloxacin concentration ratio, i.e. C_amp_/C_cipro_=115.66. The fits with the best values of α for each concentration are shown in Figure 3. The values of α are shown in Table 2. The antibiotic combination therefore appears to be more effective than the independent sum of the two single drug effects, especially at higher concentrations.

**Table 2.** Values of α with different drug combinations. α > 1 implies a synergy, whilst α < 1 implies an antagonism.

| Ciprofloxacin (mg/mL) | Ampicillin (mg/mL) | Best fit value for $\alpha$ |
| --- | --- | --- |
| 0.003 | 0.35 | $\alpha = 1.0$ |
| 0.0075 | 0.87 | $\alpha = 1.9$ |
| 0.015 | 1.7 | $\alpha = 2.4$ |
| 0.037 | 4.3 | $\alpha = 2.4$ |
| 0.075 | 8.7 | $\alpha = 2.7$ |
| 0.15 | 17.4 | $\alpha = 2.7$ |

We attempt to fit the α values with a function to extrapolate the general behaviour of α as a function of drug concentration (Figure 4). In our range of concentrations, we found a power function 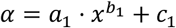 as the best fit, where x is the drug concentration. However, this function is entirely empirical. For the sake of simplicity, we only reported the ciprofloxacin concentration on the x-axis of Figure 4; the ampicillin concentration is proportional to these values (i.e. same multiples of MICs).

**Figure 4.**
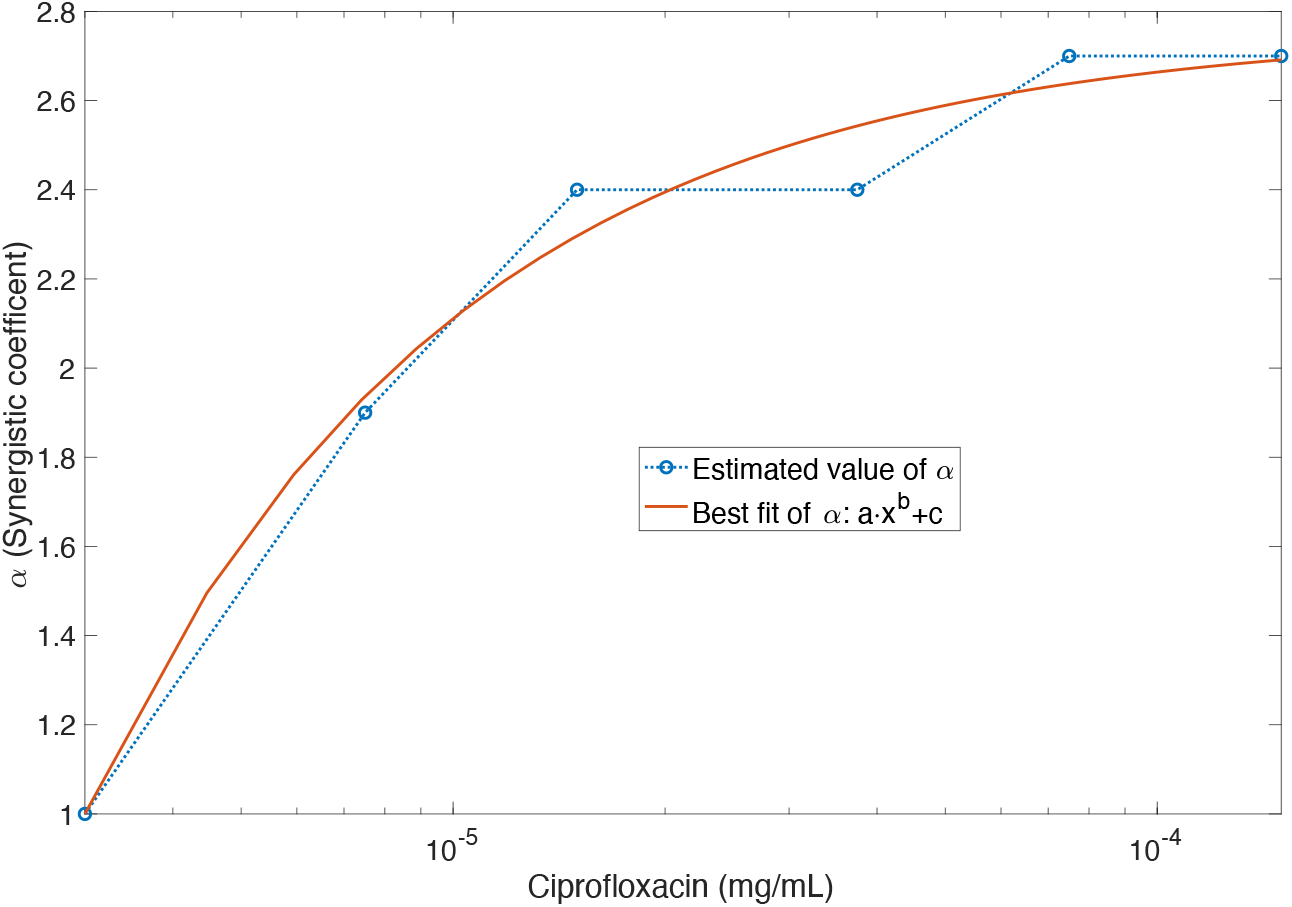
The fit of the estimated values of α. *We used a power function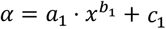, with an RMSE=0.1103, Adjusted R-square: 0.9712, and*: *a*_1_ = −4.645 ⋅ 10^−5^ (−0.0004031, 0.0003102); *b*_1_ = −0.8291 (−1.423, −0.2348); *c*_1_ = 2.76 (2.368, 3.151), ∀x > 0.

#### 3.1.3 Simulations and comparison with monotherapies

Here, we aim to evaluate the actual efficacy of the ampicillin-ciprofloxacin combination by comparing it with the ciprofloxacin and ampicillin monotherapies.

We calculate the net growth rate of the cell population treated with the drug combination. The net growth rate of the ciprofloxacin-ampicillin combination compared with ciprofloxacin monotherapy gives us the result shown in Figure 5A. In Figure 5B, there is a comparison between the net growth rate of the combination and the net growth rate of ampicillin monotherapy.

**Figure 5.**
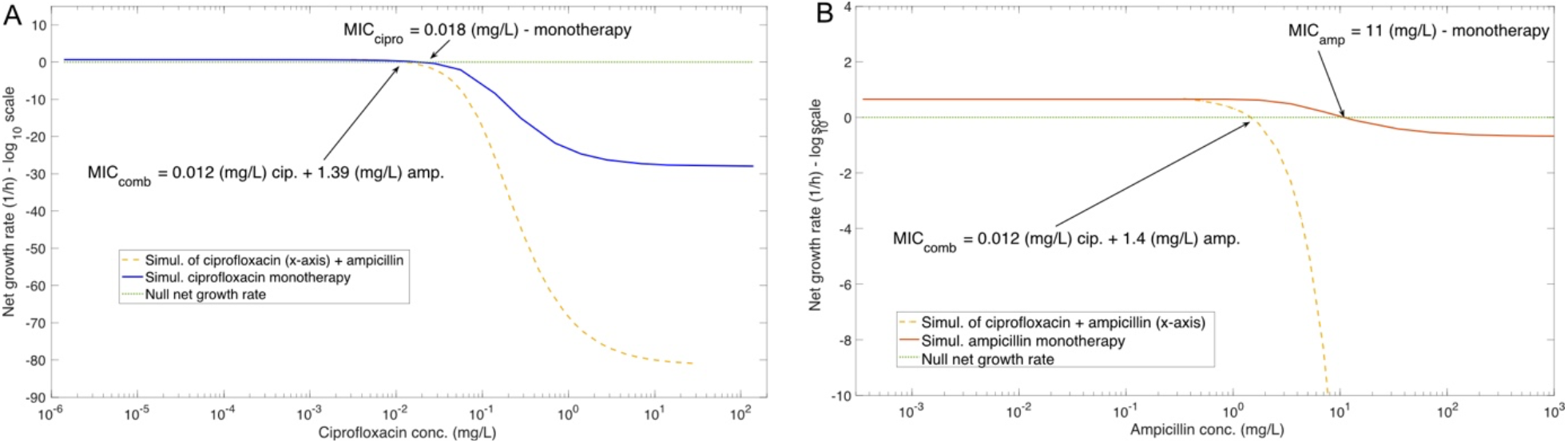
**A**: The ciprofloxacin concentration is on the x-axis. The net growth rate of ciprofloxacin is the blue curve. The yellow curve represents the net growth rate of the drug combination, where the amount of ciprofloxacin in the combination is the value on the x-axis, the value of ampicillin in the combination (not on the axis) is the concentration (ciprofloxacin)*115.66. The addition of ampicillin slightly improves the effect of ciprofloxacin. **B:** The ampicillin concentration is on the x-axis. The net growth rate of ampicillin is the red curve. The yellow curve represents the net growth rate of the drug combination, where the concentration of ampicillin in the combination is the value on the x-axis, the value of ciprofloxacin in the combination (not on the axis) is the concentration (ampicillin)/115.66. The addition of ciprofloxacin substantially improves the effect of ampicillin monotherapy, while the addition of ampicillin slightly improves the effects of ciprofloxacin monotherapy.

### 3.2 Discussion of possible interactions between ampicillin and ciprofloxacin

We observe a synergistic effect (value of α) if we compare the actual combination of ampicillin-ciprofloxacin with the independent (theoretical) combination; the synergy is more pronounced at high concentrations (Table 2). We observe a slightly improved efficacy when we compare the combination with ciprofloxacin monotherapy (Figure 5A), and a substantially improved effect when we compare the combination with the ampicillin monotherapy (Figure 5B). In Figures 5A and 5B, we observe a reduced MIC, with the drug combination indicating a more efficient action. We also note that the combination is more effective in killing the bacteria (minimum of the net growth rates in Figures 5A and 5B).

We introduced the coefficient α (Figure 4) to quantify the synergistic or antagonistic effect, but it provides no insights into the underlying mechanistic biology. Although we do not know the actual mechanism(s) of the interactions between ampicillin and ciprofloxacin, we can use our model to generate some simple hypotheses to test the flexibility of our model.

Instead of using the empirical parameter α, we can also incorporate explicit molecular mechanisms of drug-drug interactions. For example, one can model that the effective concentration of one drug inside the cell increases in the presence of another drug. As the action of ampicillin inhibits the final stages of cell wall synthesis and weakens the ability of bacteria to keep drugs outside their walls, we can assume that combination therapy leads to a scenario where the intracellular concentration of ciprofloxacin will be higher than with monotherapy.

#### 3.2.1 Hypothesis 1: Higher intracellular ciprofloxacin concentration

We now estimate the additional ciprofloxacin inside of the cells. Previously, we fitted the killing curves data (combination) with our model containing α as a free parameter in the death rate. Now, instead of using α (we set α = 1) to explain the synergy, we assume an equivalent problem with an increased ciprofloxacin concentration in the presence of ampicillin, and estimate a multiplicative free parameter by fitting the experimental killing curves.

The best fit is in Figure 6A. Figure 6B compares the previous ciprofloxacin concentration and the current higher ciprofloxacin concentration, which fits the experimental data. We observe how the ratio between the two ciprofloxacin concentrations grows at a higher concentration.

**Figure 6.**
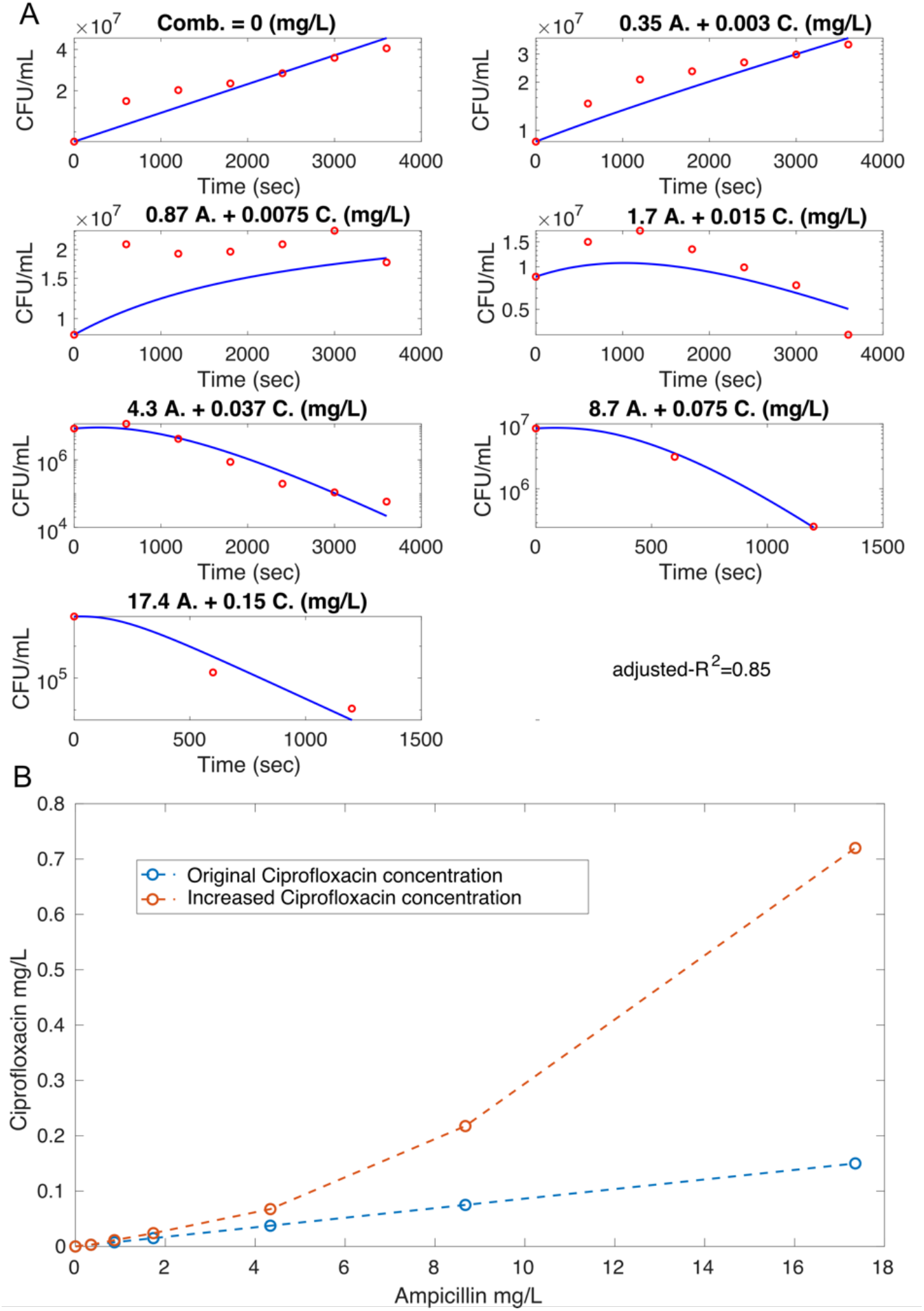
**A:** Fit with increased concentration of ciprofloxacin instead of using α to explain the synergistic effect. **B**: Comparison between the initial ciprofloxacin concentration and the increased ciprofloxacin concentrations. The ratio between the initial and the increased ciprofloxacin concentration is: (1, 1.5, 1.6, 1.8, 2.9, 4.8). The first point, zero (no drug), is not on this list.

#### 3.2.2 Hypothesis 2: Effect of increased ciprofloxacin concentration

Alternatively, the effective intracellular concentration of ciprofloxacin could be the same in both absence and presence of ampicillin. Instead, each target molecule bound by ciprofloxacin could have a stronger effect on bacterial replication and death in the presence of ampicillin.

We fit the experimental data obtained with the combination of ampicillin and ciprofloxacin, using ciprofloxacin’s growth and death rates as free parameters. We obtain the best fit in Figure 7A. Figure 7B illustrates the new growth rate obtained from the best fit in Figure 7A, and Figure 7C illustrates the new death rate.

**Figure 7.**
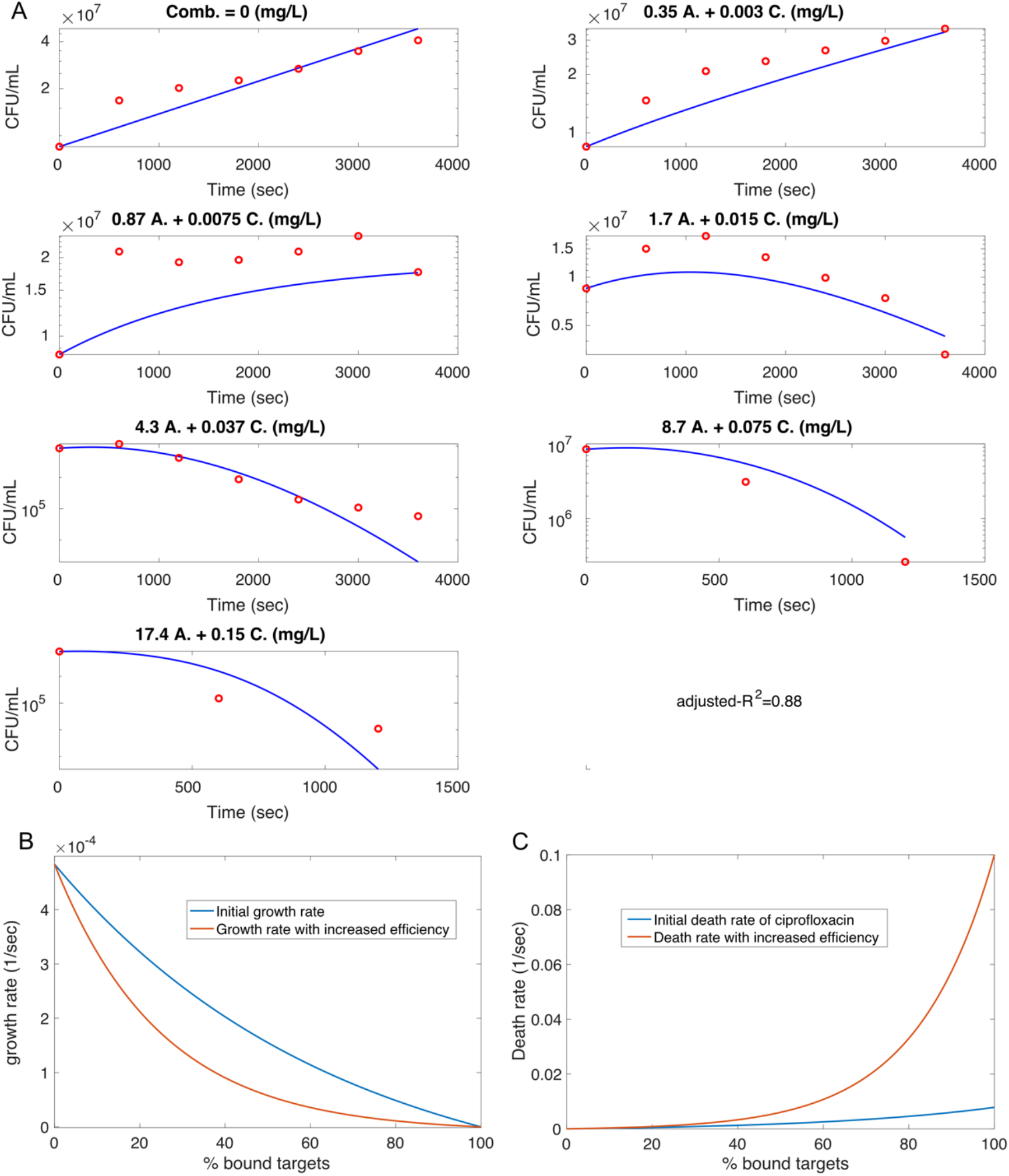
**A:** In these figures, we have our model’s best fit with the ampicillin-ciprofloxacin combination’s time-kill curves. The result of the fit gives us the adjusted growth and death rate of ciprofloxacin. **B:** Comparison between ciprofloxacin’s more efficient growth rates (in red) and the initial growth rate (in blue). **C:** Comparison between ciprofloxacin’s more efficient death rate (in red) and the initial death rate (in blue).

## 4. Discussion

We propose a mathematical model to quantify synergistic and antagonistic effects of multidrug treatment and estimate optimal drug combinations. We quantify the response of the bacterial population when exposed to drug combinations (exposure-response), and, at the same time, we can investigate different hypotheses as causes of the multidrug combined effects. We propose a novel definition of independent drug action based on a mechanistic model. We describe the drugs’ effects as a direct consequence of the binding of drug molecules to their targets, which affects the growth and death rates of bacteria. We do not use classical Hill curve or Emax models [18, 19]; but use mechanistic drug-target binding models [15, 17]. Previous models have only connected drug-target binding to bacterial death, but not bacterial replication. Such an approach is appropriate for antimicrobial peptides [13] and purely bactericidal antibiotics, but not for antibiotics that inhibit replication, i.e. those with at least partial bacteriostatic action.

Our model extends the available tools to simulate and predict drug effects, i.e. pharmacodynamics. The importance of bottom-up models in drug development is increasing. When the role of the model in drug development has a medium-high impact (e.g. data extrapolation, extending use to special populations), then a mechanistic approach becomes more reliable than a descriptive one [25, 26].

We tested our model with experimental data of *E. coli* exposed to a previously published [19] combination of ciprofloxacin and ampicillin. We proposed possible simple interpretations of the observed effects of drug combinations to highlight some advantages of using a mechanistic approach over a descriptive approach. Of note, we find that the two drugs are synergistic, especially at high concentrations, which could theoretically be explained by an increased intracellular ciprofloxacin concentration because ampicillin disrupts the cell envelope. Previously, it was only noted with this dataset that both drugs given at equal potency (each 0.5 MIC) are more effective than 1 MIC ampicillin and less effective than 1 MIC ciprofloxacin [19].

In principle, our model can be used to evaluate the effect of any combination of two drugs. What is required is empirical data to enable a matrix setup, with concentration ranges of one drug in the columns and the other in the rows. In addition, the model can facilitate the generation and testing of theoretical hypotheses about drug combinations. Finally, our model can be used to determine the likely effect of a specific molecular interaction, for example, a prediction of increased efficacy when drug penetration is increased in the presence of a second drug.

Since some treatment regimens require more than two drugs, we show how to extend our model to four drugs. A four-drug model works exactly like the model for two drugs, with the two-dimensional matrix for drug concentration expanded to a four-dimensional tensor. The simple development of a four-drug model highlights the flexibility of our approach.

The introduction of a model that is capable of predicting the efficacy of multidrug treatments can be a helpful tool in fighting wild-type, resistant, and multidrug-resistant bacterial infections. In addition to facilitating a better understanding of antibiotic action and interaction mechanisms, proposing a new mechanistic pharmacodynamic approach, as we do here, responds to the existing needs of clinical, public health and drug development agencies.

## Acknowledgments

This work is dedicated to the memory of Howie Weiss, who died a few months before the first submission. After his last comments, the draft was shortened and some language was changed, but no new simulations or interpretations were added. This work was funded by a Horizon 2020 consortium grant (anTBiotic, 733079) to PAzW. We are grateful to Sören Abel, Giovanni Montani and Angelo Vannozzi for the valuable discussions and help during the development of this work. We are also grateful to Glyn Winter for language editing. A special thank you for everything to Luciano Giannini.

## Contributions

**Conceptualization**: P. Abel zur Wiesch, F. Clarelli.

**Development of the Model**: F. Clarelli, H. Weiss.

**Software and numerical methods**: F. Clarelli.

**Validation**: F. Clarelli, P. Ankomah, H. Weiss, J. M. Conway, G. Forsdhal, P. Abel zur Wiesch.

**Experimental data**: P. Ankomah.

**Writing - Original Draft**: F. Clarelli.

**Writing - Review & Editing**: F. Clarelli, P. Ankomah, H. Weiss, G. Forsdhal, J. M. Conway, P. Abel zur Wiesch.

**Project administration**: G. Forsdahl, P. Abel zur Wiesch.

**Funding acquisition**: P. Abel zur Wiesch.

