## Supplemental for "A Mechanistic Approach to Optimize Combination Antibiotic Therapy"

### Supplementary material

#### S.1 Parameter values

Modified from Table S1 in [1].

| Drug | Parameter | Value | References |
|---|---|---|---|
| Ciprofloxacin | $k_f$ | $3.5 \cdot 10^3 \ (M^{-1} \cdot sec^{-1})$ | [1,2,3] |
| | $k_r$ | $3 \cdot 10^{-4} \ (sec^{-1})$ | [1,2] |
| | # targets per bacterium | $100 \ (\#)$ | [1,4,5] |
| Ampicillin | $k_f$ | $495.7 \ (M^{-1} \cdot sec^{-1})$ | [1,6,7] |
| | $k_r$ | $1 \cdot 10^{-4} \ (sec^{-1})$ | [1,7] |
| | # targets per bacterium | $2500 \ (\#)$ | [1,8,9] |

### S.2 Single drug fit

_Ciprofloxacin_

We fit our model with ciprofloxacin time-kill curves (with seven different drug concentrations), using the coefficients of growth and death rates of bacteria as free parameters, and use the values of association and dissociation rates from the literature [1-3]. See Table 1 in section S1 of supplementary material for all the parameter values.

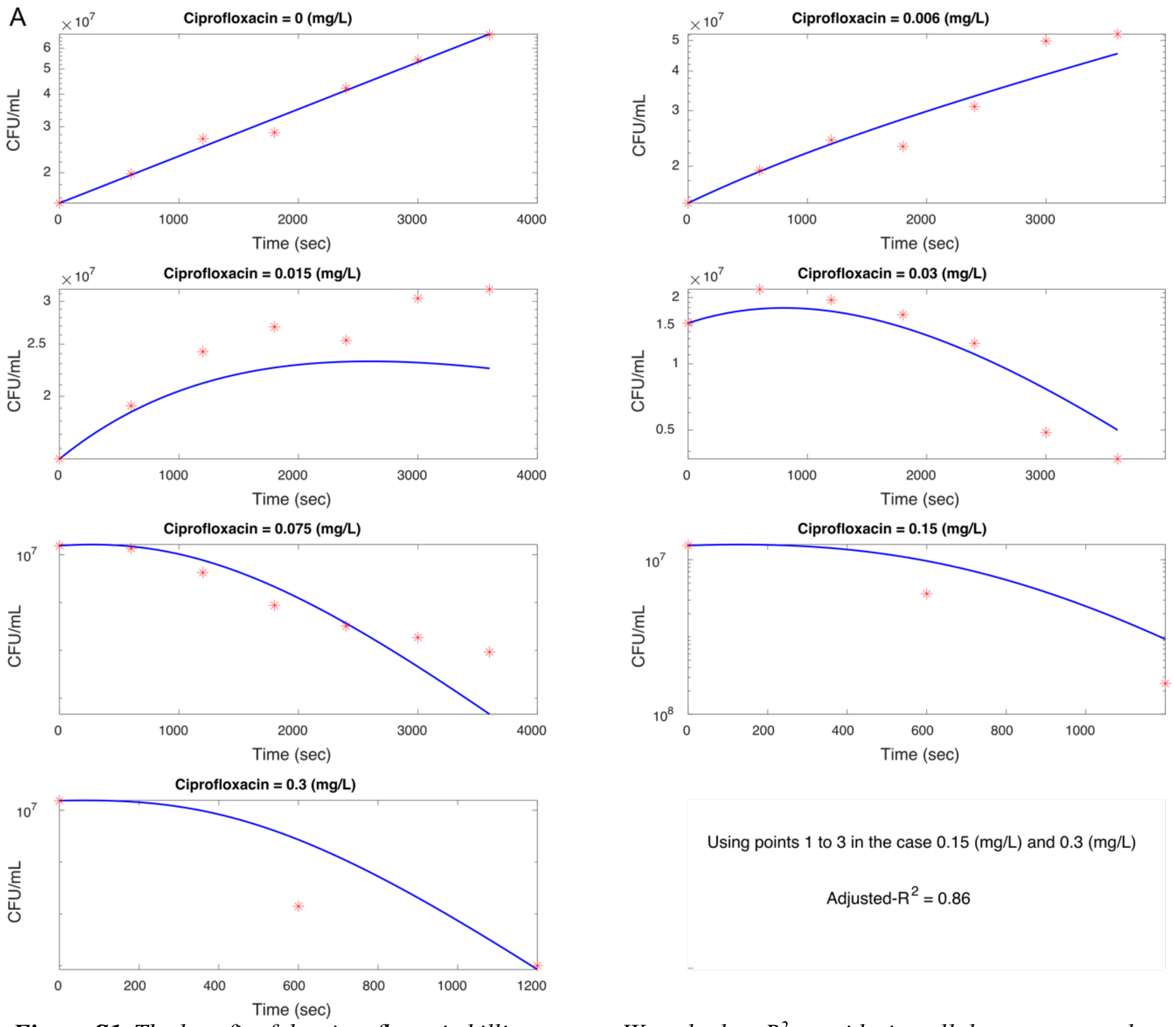


***Figure S1***. *The best fit of the ciprofloxacin killing curves. We calculate* $R^2$ *considering all the curves together. We consider only the first three points in the third plot in the right-hand column (ciprofloxacin = 0.15 mg/l) and the fourth plot in the left-hand column (ciprofloxacin = 0.3 mg/l) because of the persistence appearing in the successive points.*

The fit in Figure S1 has been realized in different steps. The maximum growth rate has been estimated fitting the growth of bacteria without drugs. This value can be slightly different for the ciprofloxacin monotherapy, ampicillin monotherapy and in the combination of the two drugs.

The maximum death rate for bacteria exposed to ciprofloxacin is estimated with the highest drug concentration. In this case, the highest experimental concentration (0.3 mg/L) is not high enough to give us the maximum death rate for our model. Considering the association and dissociation rate we are using, with a ciprofloxacin concentration equal to 0.3 mg/L, we find that the maximum number of bound targets is around 91% of the total.

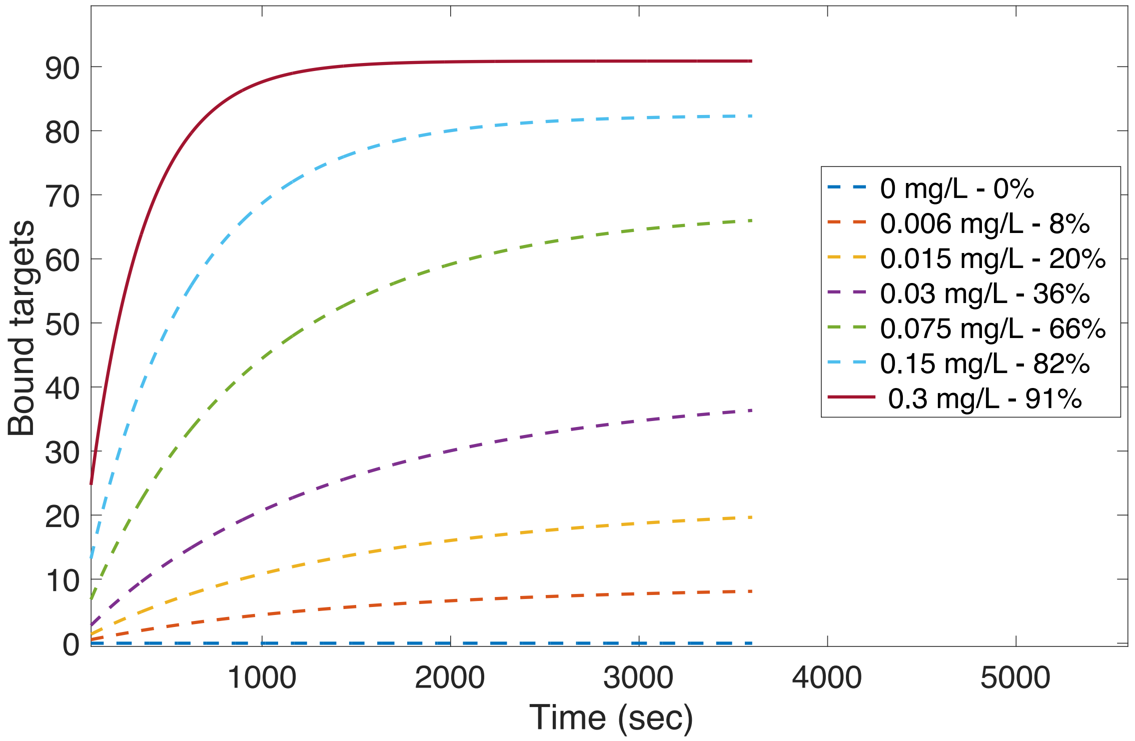


***Figure S2***. *The number of bound targets reached with each ciprofloxacin concentration used in the experiments. The highest concentration gives us 91% of the maximum number of bound targets.*

We need a maximum death rate that gives us a percentage of bound targets close to 100% of the total bound targets.

The maximum death rate we obtain fitting the kill curve with 0.3 mg/L of ciprofloxacin is 0.0078 (1/sec). If we use this value as the maximum death rate, we obtain the simulation in Figure S2, which is smaller than what we need, based on our model requirements.

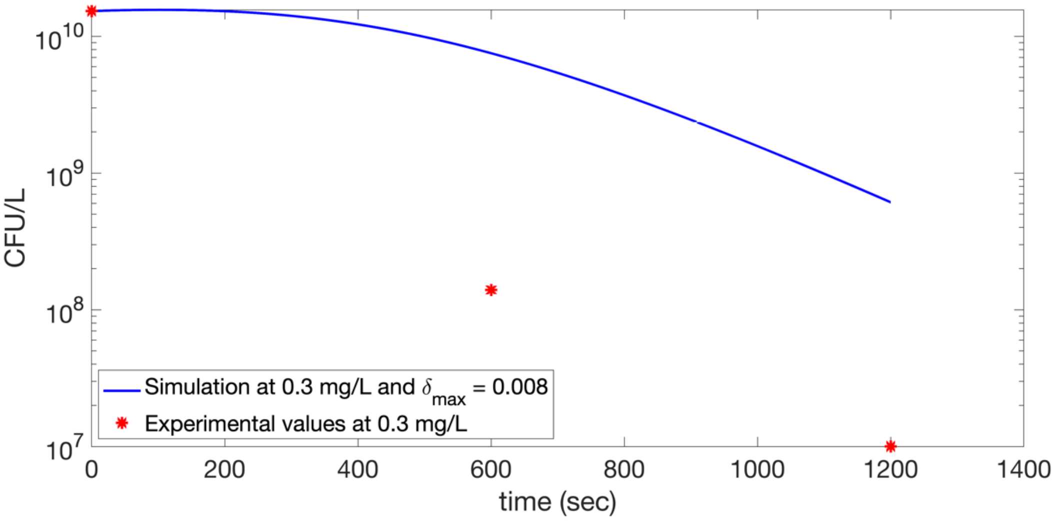


***Figure S3***. *Simulation with 0.3 mg/L of ciprofloxacin, using a maximum death rate* $\delta_{max} = 0.008$.

To avoid this underestimated value for the maximum growth rate, we can rescale the maximum death rate in a way that fits the experimental points shown in Figure S2. The rescaled maximum death rate is $\delta_{max}^{cipro} = 0.018\ (1/sec)$.

Once we have calibrated the maximum death rate for ciprofloxacin, the fit of the other experimental data can be done using all the concentrations at the same time, using the minimum squared error as the value to minimize. The method used in the optimization is the particle swarm optimization (PSO), whilst the numerical schemes to solve the differential equations are developed using a finite difference scheme.

The growth and death rates in Figure S4A are the rates for the best fit.

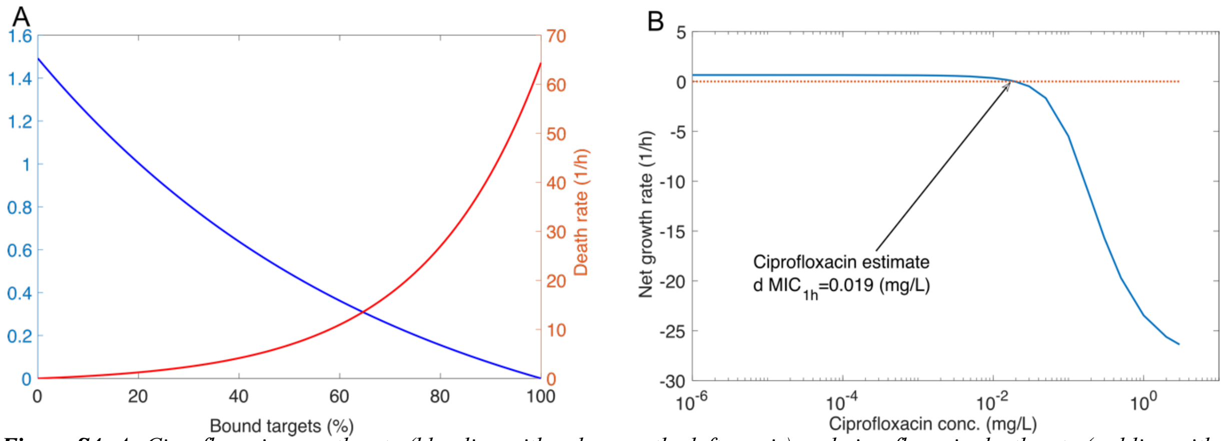


***Figure S4. A:*** *Ciprofloxacin growth rate (blue line with values on the left y-axis) and ciprofloxacin death rate (red line with values on the right y-axis) as a function of bound targets. We obtain these functions by the best fit of the killing curves in Figure 3A.* ***B***: *Simulations of the bacterial response under different ciprofloxacin concentrations. The y-axis represents the net growth rate, i.e.* $\left(log(B(1h)) - log(B(0))\right)/1h$, *as a function of ciprofloxacin concentration. In this relationship, B represents the number of bacteria. We estimate the value of 0.019 (mg/L) for the MIC.*

With our model calibrated with ciprofloxacin monotherapy, we simulate the bacterial response under a wide range of drug concentrations. We plot the results of these simulations in Figure 3B, where the net growth rate of bacteria is a function of the drug concentration. We define as net growth rate the difference $\frac{\log(B(t_{fin}))-\log(B(t_0))}{t_{fin}-t_0}$. We estimate the minimum inhibitory concentration (MIC) to be when the net growth rate intersects the null growth line – i.e. when $B(t_{fin}) = B(t_0)$ (see Figure S4B).

*Ampicillin*

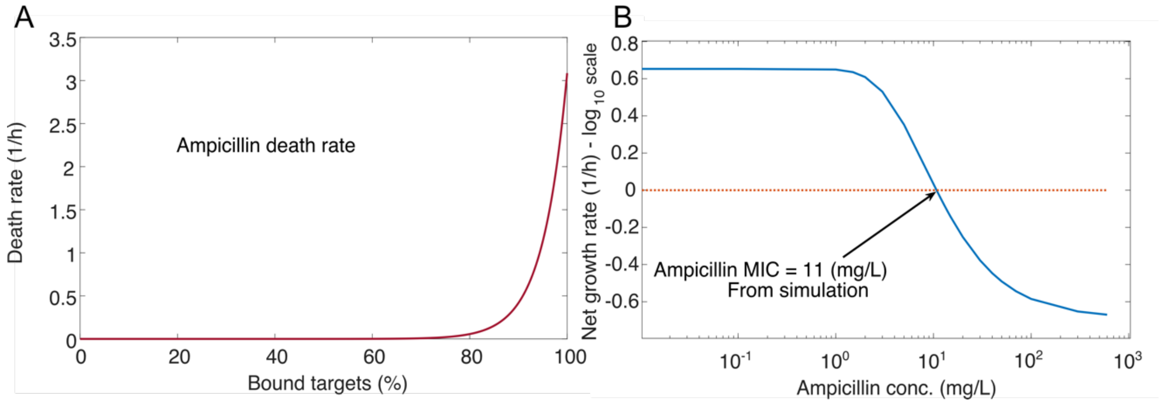


***Figure S5. A***: *The red curve is the ampicillin death rate as a function of bound targets. This result comes from the best fit presented in Fig. A.* ***B***: *Simulations of the bacterial response under different concentrations of ampicillin. On the y-axis, the net growth rate of bacteria, i.e.* $(log(B(1h)) - log(B(0)))/1h$, *as a function of ampicillin concentration. B represents the number of bacteria. We estimate the value of 11 (mg/L) for the MIC.*

This section repeats the previous analysis but for ampicillin. However, the action of ampicillin can be considered mainly bactericidal. Therefore, ampicillin's minimal bacteriostatic activity makes the effect on growth rate negligible, so the model fit provides only a death rate (as shown by figure S5A, where only the bacterial death rate depends on the antibiotics). In addition, as for ciprofloxacin, we use values from the literature for the association and dissociation rates (see table in the supplementary material). We used the same approach to estimate the maximum growth rate in as for ciprofloxacin (described above). In this case, we obtain the rescaled value of $\delta_{max}^{amp} = 0.00086\ (1/sec)$.

The best fit of our model with the ampicillin time-kill curves is shown in Figure S6 (supplementary material), and, based on this result, we obtain the ampicillin death rate in Figure S5A. After the calibration, we simulate the bacterial response under a wide range of ampicillin concentrations, with the results shown in Figure S5B. The estimated ampicillin MIC is 11 mg/L, which is represented by the intersection with the null dotted line in Figure S5B.

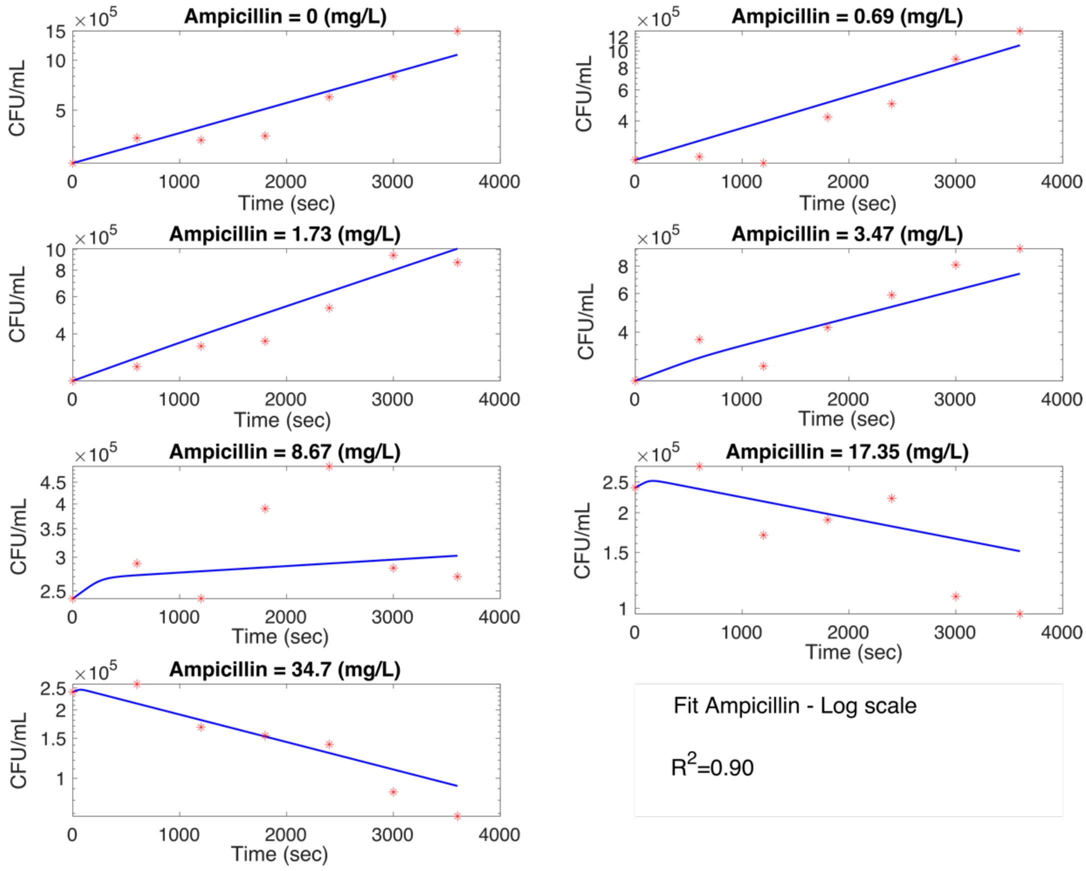


***Figure S6.*** *The best fit of the ampicillin killing curves. We also calculated the adjusted* $R^2$ *using all the curves fitted together.*
